## Supplementary material for "HetL provides immunity to HetR against PatS inhibition, and promotes pattern formation in the cyanobacterium *Nostoc* PCC 7120": HetL supplemental data

#### Supplementary Table 1: PRPs encoding genes in the genome of *Nostoc*

| Gene ID/name | Num of AA | Num of PRs domains | Additional Domain(s) | Cellular location | Gene expression (number of reads) | | | |
|---|---|---|---|---|---|---|---|---|
| | | | | | 0h | 6h | 12h | 21h |
| **Genes whose expression changes in response to combined nitrogen starvation** | | | | | | | | |
| alr4610 | 164 | 2 | - | P/L | 33 | 30 | 88 | 92 |
| alr1298 | 167 | 1 | - | C | 79 | 65 | 151 | 119 |
| all0186 | 168 | 2 | - | M | 13 | 6 | 74 | 30 |
| alr1746 | 182 | 3 | - | C | 843 | 519 | 134 | 159 |
| all4303 | 213 | 4 | - | C | 39 | 51 | 33 | 80 |
| all3048 | 217 | 2 | DnaJ | C | 96 | 84 | 131 | 195 |
| all2395 (FraF) | 222 | 3 | - | C | 15 | 11 | 105 | 80 |
| alr1579 | 222 | 3 | - | P/L | 4 | 5 | 67 | 8 |
| all3740 (HetL) | 237 | 4 | - | C | 16 | 18 | 50 | 78 |
| all4152 | 450 | 1 | - | M | 9 | 21 | 28 | 27 |
| all3305(PatL) | 496 | 5 | - | M | 76 | 94 | 163 | 137 |
| alr3268 | 524 | 2 | Kinase | C | 6 | 8 | 17 | 23 |
| all3114 | 576 | 5 | - | C | 19 | 17 | 56 | 68 |
| alr9014* | 679 | 3 | - | M | 40 | 40 | 87 | 73 |
| alr0704 | 693 | 3 | - | M | 21 | 28 | 39 | 42 |
| all0813 (HglK) | 727 | 4 | RDD domain | M | 65 | 52 | 316 | 222 |
| **Genes whose expression is not impacted by combined nitrogen starvation** | | | | | | | | |
| all1812 | 125 | 1 | - | C | | | | |
| alr5209 | 129 | 2 | - | C | | | | |
| alr0433 | 143 | 2 | - | P/L | | | | |
| all4220 | 152 | 1 | - | P/L | | | | |
| alr2741 | 182 | 2 | - | M | | | | |
| all3332 | 206 | 1 | | M | | | | |
| all3306 | 252 | 2 | - | C | | | | |
| alr2768 | 256 | 1 | - | L/P | | | | |
| all3256 | 268 | 4 | - | C | | | | |
| alr7125* | 369 | 2 | - | C | | | | |
| all3869 | 376 | 4 | Endoribonuclease L-PSP | C | | | | |
| all0958 | 475 | 2 | - | C | | | | |
| alr1142 | 521 | 3 | Pentapeptide 4 (9PRs) | C | | | | |
| alr7124* | 586 | 2 | - | M | | | | |
| alr3131 | 953 | 2 | - | C | | | | |
| all8023 | 1010 | 3 | - | C | | | | |

The sequences were retrieved from the Microscope Mage database. The genes located on *Nostoc* plasmids are marked by an asterisk. The functional domains were analyzed by Pfam. The cellular localization was deduced from the presence of transmembrane domain, signal peptide and lipopeptide using TMhmm, SignalIP and LipoP softwires. For gene expression, the reads values are those reported in {Flaherty, 2011 #352}.

#### Supplementary table 2

#### List of active residues used as input for HetR-Hood :HetL docking simulations

| ***Protein*** | ***Active residues number*** |
|---|---|
| HetL | 0,1,2,3,5,7,8,9,10,11,12,13,14,15,16,18,19,20,21,23,24,26,28,29,31,33,36,39,43,44,46,49,51,53,54,58,59,61,64,66,68,71,74,78,79,81,83,84,88,89,91,93,94,96,98,99,103,104,106,108,111,113,114,116,119,121,123,124,126,128,129,130,131,132,133,134,136,139,141,143,144,146,149,151,153,158,159,161,163,164,166,169,171,174,175,176,177,178,179,180,181,184,186,187,189,191,192,194,197,199,201,204,206,207,209,211,212,214,215,216,217,218,219,220,221,222,224,225,226,227,228,230,231,232,233,234,235,236 |
| HetR-Hood | 222,223,224,225,227,228,229,231,233,235,236,239,240,241,243,244,245,246,247,249,250,252,253,254,256,259,260,261,262,263,264,266,267,268,269,271,272,273,274,276,279,280,281,282,283,285,287,296,297,298 |

### Supplementary file 1:

#### *Nostoc* and *E. coli* strains

| Strains | Description/genotype | Source/reference |
|---|---|---|
| ***Nostoc* strains** | | |
| *Nostoc* PCC 7120 | Wild type strain (WT) | Pasteur Institute Collection |
| WT/P*petE-patS* | WT strain containing the pRL1272-P*petE-patS* plasmid | This study |
| WT/P*petE-patS* P*petE-hetL* | WT strain containing the pRL1272-P*petE-patS* plasmid and pRL25T-P*petE-hetL* plasmid | This study |
| WT/P*petE-patS* P*petE-hetL*D151A | WT strain containing the pRL1272-P*petE-patS* plasmid and pRL25T-P*petE-hetL*D151A plasmid derivative where the *hetL* gene has been mutated to encode for a D151A substitution | This study |
| WT/P*petE-hetL* | WT strain containing the pRL25T-P*petE-hetL* plasmid | This study |
| WT/P*petE-hetL*D151A | WT strain containing the pRL25T-P*petE-hetL*D151A plasmid | This study |
| WT/P*patS-hetL* | WT strain containing the pRL25T-P*patS-hetL* plasmid | This study |
| WT/P*rbcL-hetL* | WT strain containing the pRL25T-P*rbcL-hetL* plasmid | This study |
| Δ*hetR* | *Nostoc* deletion mutant of the *hetR* gene | Borthakur et al., 2005 |
| ***E. coli* strains** | | |
| TG1 | *supE thi*-1 Δ(*lac-proAB*) Δ(*mcrB-hsdSM*)5 (rK– mK–) [F´ *traD36 proAB lacIqZΔM15*] | Stratagene |
| DH5α | *fhuA2 lac(del)U169 phoA glnV44 Φ80' lacZ(del)M15 gyrA96 recA1 relA1 endA1 thi-1 hsdR17* | Taylor, RG et al. (1993) |
| Stellar™ | *F–, endA1, supE44, thi-1, recA1, relA1, gyrA96, phoA, Φ80d lacZΔ M15, Δ (lacZYA - argF) U169, Δ (mrr - hsdRMS - mcrBC), ΔmcrA, λ–* | Takara |
| BL21 DE3 | *fhuA2 [lon] ompT gal (λ DE3) [dcm] ΔhsdS λDE3 = λsBamHIo ΔEcoRI-int::(lacI::PlacUV5::T7 gene1) i21Δnin5* | NEB |
| BTH101 | $F^{-}$*, cya-99, araD139, galE15, galK16, rpsL1, hsdR2, mcrA1, mcrB1* | Karimova G et al, 1998 |

| | | |
|---|---|---|
| eXX1 | TG1 containing the plasmid pKT25-*hetR* | This study |
| eXX2 | TG1 containing the plasmid pKT25-*hetL* | This study |
| eXX3 | TG1 containing the plasmid pKT25-*hetL*D151A | This study |
| eXX4 | TG1 containing the plasmid pUT18C-*hetR* | This study |
| eXX5 | TG1 containing the plasmid pUT18C-*hetR*R223W | This study |
| eXX6 | TG1 containing the plasmid pUT18C-*hetL* | This study |
| eXX7 | TG1 containing the plasmid pUT18C-*hetL* RBS-*patS* | This study |
| eXX8 | TG1 containing the plasmid *hetR*$_{\text{hood}}$-pUT18 | This study |
| eXX9 | TG1 containing the plasmid pUT18C-*all3256* | This study |
| eXX10 | TG1 containing the plasmid pUT18C-*all4303* | This study |
| eBR1 | TG1 containing the plasmid pET28a-*his-hetR* | Roumezi et al., 2020 |
| eXX11 | Stellar$^{\text{TM}}$ containing the plasmid pRL1272-P*petE-patS* | This study |
| eXX12 | Stellar$^{\text{TM}}$ containing the plasmid pRL25T-P*petE-hetL* | This study |
| eXX13 | Stellar$^{\text{TM}}$ containing the plasmid pRL25T-P*petE-hetL*D151A | This study |
| eSC1 | Stellar$^{\text{TM}}$ containing the plasmid pRL25T-P*patS-hetL* | This study |
| eSC2 | Stellar$^{\text{TM}}$ containing the plasmid pRL25T-P*rbcL-hetL* | This study |
| eXX14 | BTH101 containing plasmids pKT25-zip and pUT18C-zip | This study |
| eXX15 | BTH101 containing plasmids pKT25 and pUT18C | This study |
| eXX16 | BTH101 containing plasmids pKT25-*hetL* and pUT18C | This study |
| eXX17 | BTH101 containing plasmids pKT25 and pUT18C-*hetR* | This study |
| eXX18 | BTH101 containing plasmids pKT25-*hetR* and pUT18C-*hetR* | This study |
| eXX19 | BTH101 containing plasmids pKT25-*hetL* and pUT18C-*hetR* | This study |
| eXX20 | BTH101 containing plasmids pKT25-*hetL* and pUT18C-*hetL* | This study |
| eXX21 | BTH101 containing plasmids pKT25-*hetL* and *hetR*$_{\text{hood}}$-pUT18 | This study |

| | | |
|---|---|---|
| eXX22 | BTH101 containing plasmids pKT25-*hetL*D151A and pUT18C-*hetL* | This study |
| eXX23 | BTH101 containing plasmids pKT25-*hetR* and pUT18C-*hetR*R223W | This study |
| eXX24 | BTH101 containing plasmids pKT25-*hetL*D151A and pUT18C-*hetR* | This study |
| eXX25 | BTH101 containing plasmids pKT25-*hetL* and pUT18C-*hetR*R223W | This study |
| eXX26 | BTH101 containing plasmids pKT25-*hetR* and pUT18C-*hetL* | This study |
| eXX27 | BTH101 containing plasmids pKT25-*hetR* and pUT18C-*hetL*-RBS-*patS* | This study |
| eXX28 | BTH101 containing plasmids pKT25-*hetR* and pUT18C-*hetL*-RBS-*patS6* | This study |
| eXX29 | BTH101 containing plasmids pKT25-*hetR* and pUT18C-*all3256* | This study |
| eXX30 | BTH101 containing plasmids pKT25-*hetR* and pUT18C-*all4303* | This study |

**Plasmids**

| **Plasmids** | **Description** | **Source/reference** |
|---|---|---|
| pKT25-zip | Two hybrid plasmid Kan$^{R}$ | {Karimova, 1998 #130} |
| pUT18C-zip | Two hybrid plasmid Amp$^{R}$ | {Karimova, 1998 #130} |
| pKT25 | Two hybrid plasmid. T25 at the N terminus Kan$^{R}$ | {Karimova, 1998 #130} |
| pUT18C | Two hybrid plasmid. T18 at the N terminus Amp$^{R}$ | {Karimova, 1998 #130} |
| pKNT25 | Two hybrid plasmid. T25 at the C terminus Kan$^{R}$ | {Karimova, 1998 #130} |
| pUT18 | Two hybrid plasmid. T18 at the C terminus Amp$^{R}$ | {Karimova, 1998 #130} |
| pET28a | His-tagged protein expression plasmid in *E. Coli* Kan$^{R}$ | Novagen |
| pRL1272 | Replicative in *Nostoc* Ery$^{R}$ | {Wolk, 1988 #308} |
| pRL25T | Replicative in *Nostoc* Neo$^{R}$ | {Yang, 2013 #307} |
| pXX1 | pKT25-*hetR* | This study |
| pXX2 | pKT25-*hetL* | This study |
| pXX3 | pKT25-*hetL*D151A | This study |
| pXX4 | pUT18C-*hetR* | This study |
| pXX5 | pUT18C-*hetR*R223W | This study |
| pXX6 | pUT18C-*hetL* | This study |
| pXX7 | pUT18C-*hetL*-RBS-*patS* | This study |
| pXX8 | pUT18C-*hetL*-RBS-*patS6* | This study |
| pXX9 | *hetR*$_{\text{hood}}$-pUT18 | This study |
| pXX10 | pUT18C-*all3256* | This study |
| pXX11 | pUT18C-*all4303* | This study |
| pBR1 | pET28a-*his-hetR* | Roumezi et al., 2020 |
| pXX12 | pET28a-*hetL-his* | This study |
| pCSB270 | pRL1272-P*petE* | This study |
| pXX13 | pRL1272-P*petE-patS* | This study |
| pCSB265 | pRL25T-P*petE* | This study |
| pXX14 | pRL25T-P*petE-hetL* | This study |
| pXX15 | pRL25T-P*petE-hetL*D151A | This study |
| pSC1 | pRL25T-P*patS-hetL* | This study |
| pSC2 | pRL25T-P*rbcL-hetL* | This study |

#### Primers

| Name | Sequence (5’-3’) | Experiment |
|---|---|---|
| 16S rRNA rt fw | TCCTGGTGTAGCGGTGAAAT | Quantitative RT-PCR analysis |
| 16S rRNA rt rv | AGCCACGCCTAGTATCCATC | |
| *hetP* rt fw | TGGCTGGTAAATACTCTTGGG | |
| *hetP* rt rv | ACCTACTACTTCCAGATAGGC | |
| *hetL* rt fw | GACATTATGCTGCTGGCAAA | |
| *hetL* rt rv | CAAGTCGCGTCTGACGTAAA | |
| *hetR* rt fw | GCGTCGTCTGCTTTACTCTG | |
| *hetR* rt rv | CCCAGTCTTTCATCATGCGG | |
| *hetR* dh fw T25 | TTTTCTGCAGGGATGAGTAACGACATCGATCT | |
| *hetR* dh fw T18 | TTTTCTGCAGGATGAGTAACGACATCGATCTGA | |
| *hetR* dh rv | TTTTTGAATTCTTAATCTTCTTTTCTACCAAACACCATTTG | Two hybrid assays |
| Mut *hetR*R223W fw | CCAGCAGACGACCAAGAGTGGACTTATATTATGGTGGAA | |
| Mut *hetR*R223W rv | TTCCACCATAATATAAGTCCACTCTTGGTCGTCTGCTGG | |
| *hetL* dh fw T25 | TTTTCTGCAGGGATGAATGTGGGTGAAAT | |
| *hetL* dh fw T18 | TTTTCTGCAGGATGAATGTGGGTGAAAT | |
| *hetL* dh rv | TTTTTGAATTCTCAATCATGAATTGAACCATCAGG | |
| RBS-*patS dh* fw T18 | TTTTTGTCGACAGGTTAGGAGAACCATATG | |
| RBS-*patS dh* rv T18 | TTTTTCTCGAGGATTGAGTGGTCGGAACGA | |
| RBS-*patS6 dh* fw T18 | AAGCTTATCGATACCGTCGACAGGTTAGGAGAACCATATGGAGCGCGGTAGTGGTAGATAGAACG | |
| RBS-*patS6 dh* rv T18 | GATGAATTGCTCGAGGTCGACGATTGAGTGGTCGGAACGAATGC | |
| *hetR*$_{\text{hood}}$ dh fw | TTTTCTGCAGGTATGCCCCAGCAGAC | |
| *hetR*$_{\text{hood}}$ dh rv | TTTTTGAATTCGATCTACCAAACACCATTTGTAAAATCATGG | |
| *all3256* dh fw T18 | TTTTCTGCAGGATGGCAAATCTAGAGCAT | |
| *all3256* dh rv | TTTTTGAATTCCTAATCATGCCTTGAAGAGTCA | |
| *All4303* dh fw T18 | TTTTCTGCAGGATGAATATTGACGCTATT | |
| *All4303* dh rv | TTTTTGAATTCTTACCCATTACCAATTTCTAATATTGTCCCT | |

| | | |
|---|---|---|
| Mut *hetL*D151A fw | GCAGATTTAAGCTACGCTGCCCTGAGAGCGGCTTCTCTA | |
| Mut *hetL*D151A rv | TAGAGAAGCCGCTCTCAGGGCAGCGTAGCTTAAATCTGC | |
| *hetR* pET28 fw | CATATGAGTAACGACATCGATCTG | |
| *hetR* pET28 rv | GGATCCTTAATCTTCTTTTCTACC | |
| *hetL* pET28 fw | AGGAGATATACCATGGGCAATGTGGGTGAAATTCTGAGACA | Protein production for BLI assays |
| *hetL* pET28 rv | GGTGGTGGTGCTCGAGACCTTGAAAATAAAGATTTTCATCATGAATTGAACCATCA | |
| *patS* pRL fw | GCCCATCGATGGATCCATGAAGGCAATTATGTTAGTG | |
| *patS* pRL rv | CGTCGACCCGGGATCCATGACTATTGACCAAATGACTATTG | |
| *hetL* pRL fw | GAGCTCGTCGACCCGGGATCCTCAATCATGAATTGAACCATCAGGC | Construction of recombinant plasmids for *Nostoc* |
| *hetL* pRL rv | ATGGGGCCCATCGATGGATCCATGAATGTGGGTGAAATTCTGAGAC | |
| P*patS* fw | TGAGATTATCAAAAAGGATCCAGATCCTGAATTTGTTTTGGGAAC | |
| P*patS* rv | CCACATTCATAATCTTAACCTCCCTGAATTACTTTTCAACAGAACATTT | |
| *hetL* P*patS* fw | TAAGATTATGAATGTGGGTGAAATTCTGAGAC | |
| *hetL* P*patS* rv | GAGTAGAATTCCCGGGGATCCTCAATCATGAATTGAACCATCAGGC | |
| P*rbcL* fw | TGAGATTATCAAAAAGGATCCGCAGGGGAAGTAAAGAAGAATGAC | |
| P*rbcL* rv | CACCCACATTCATATCTATCCTTCCAAGATGTCAC | |
| *hetL* P*rbcL* fw | GATAGATATGAATGTGGGTGAAATTCTGAGAC | |
| *hetL* P*rbcL* rv | GAGTAGAATTCCCGGGGATCCTCAATCATGAATTGAACCATCAGGC | |
| P*hetP* fw | [6FAM]ATTTAGTGGTAAATTCTCTT | EMSA assay |
| P*hetP* rv | TGAGTTATACGCTATATCAA | |

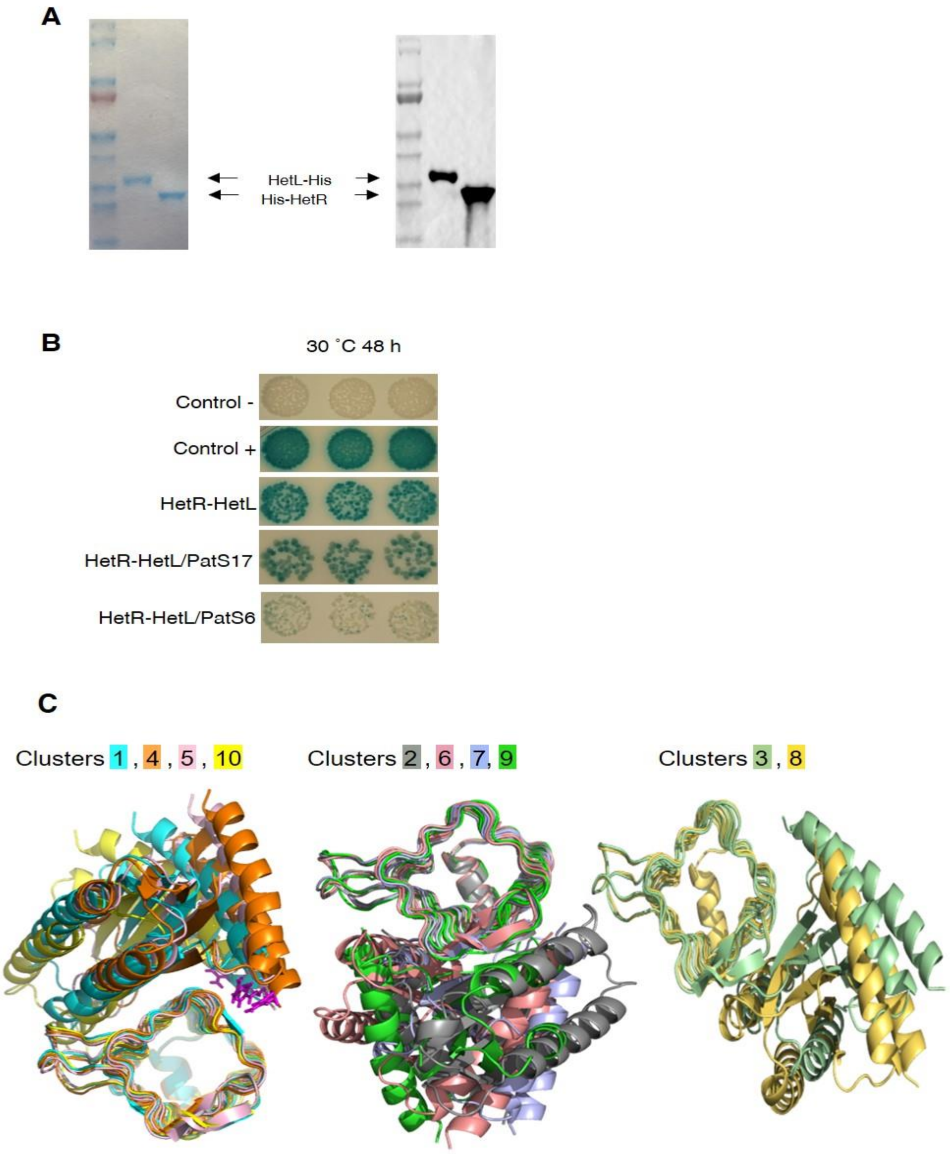

**Supplementary Figure 1.**

(**A**) Purified HetL and HetR proteins. (**left**) 1 µg purified proteins migrated in 4-20% gel (Nusep) colored by instant blue. (**right**) Anti-His immunoblot analysis of 1 µg purified proteins.

(**B**) PatS-6 interferes with HetL-HetR interaction. Bacterial two hybrid assay between HetL and HetR in the presence of PatS. Strains after transformations with indicated plasmids were grown on LB plates containing IPTG, X-gal, and corresponding antibiotics for 48 h at 30 ˚C. Strains producing the T18 and T25 served as negative control. Strains producing T18-Zip and T25-Zip served as positive control. PatS17 (the full-length PatS) and PatS6 (ERGSGR) were produced from plasmids pXX7 and pXX8 respectively.

(**C**) Best models obtained from docking simulations. Three groups composed by equivalent models are presented. Models from each group are superimposed. Each color represents a defined model. R223 residue from the best models in clusters 1, 4, 5,10 are colored in magenta.

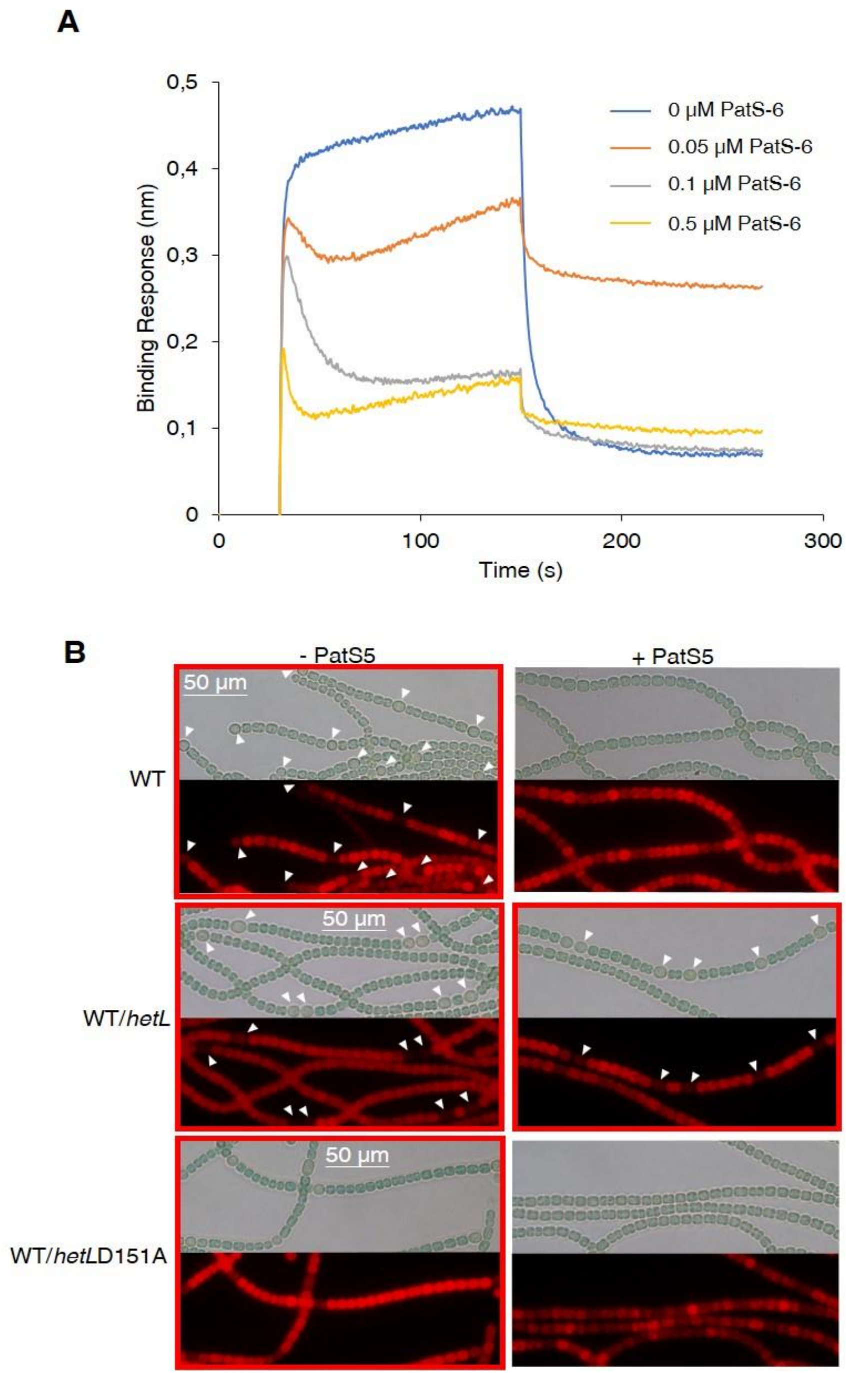

**Supplementary Figure 2.**

(**A**) BLi assay between HetL and HetR in the presence of PatS-6. 10 µM of HetL was incubated 5 mins with different concentrations of PatS-6 at 0, 0.05, 0.1 and 0.5µM before bringing HetL in contact with the bound HetR. Each curve represents the average of two experiments minus the control experiment. Control experiment HetL 10µM loaded onto a biosensor devoid of HetR.

(**B**) WT/P*petE-hetL* strain was able to form heterocysts in the presence of the PatS-5 peptide. Microscopic bright field images (upper) and auto-fluorescence images (lower) of indicated *Nostoc* strains after 24 hours of nitrogen stepdown in addition of 3 µM copper are shown. White arrows point to heterocysts. Images with heterocysts are with red frames. Similar results were obtained with PatS-6.
